# Does parental angling selection affect the behavior or metabolism of brown trout parr?

**DOI:** 10.1101/611293

**Authors:** Jenni M. Prokkola, Nico Alioravainen, Lauri Mehtätalo, Pekka Hyvärinen, Alexandre Lemopoulos, Sara Metso, Anssi Vainikka

**Author notes:** Organismal and Evolutionary Biology Research Programme, P.O. Box 56, 00014 University of Helsinki, Finland.

## Abstract

The behavior of organisms can be subject to human induced selection such as that arising from fishing. Angling is expected to induce mortality on fish with bold and explorative behavior, which are behaviors commonly linked to a high standard metabolic rate. We studied the transgenerational response of brown trout (*Salmo trutta*) to angling-induced selection by examining the behavior and metabolism of 1-year-old parr between parents that were or were not captured by experimental fly fishing. We performed the angling selection experiment on both a wild and a captive population, and compared the offspring for standard metabolic rate and behavior under predation risk in common garden conditions. Angling had population-specific effects on risk taking and exploration tendency, but no effects on standard metabolic rate. Our study adds to the evidence that angling can induce transgenerational responses on fish personality. However, understanding the mechanisms of divergent responses between the populations requires further study on the selectivity of angling in various conditions.

## Introduction

Survival selection by hunting and fishing can differ from natural selection patterns and induce phenotypic changes in various traits over time (Fugère and Hendry 2018). At worst, anthropogenic selection can increase the relative frequencies of maladaptive phenotypes decreasing the fitness of harvested populations (Allendorf and Hard 2009; Coltman et al. 2003). Experimental studies have shown that responses to human-induced selection can be rapid at both phenotypic, including behavior (Kern et al. 2016; Sbragaglia et al. 2019; Wong et al. 2012) and genetic levels (Bowles et al. 2020; Cooke et al. 2007; Sutter et al. 2012; Uusi-Heikkilä et al. 2015). Fisheries-induced selection can occur on traits that explain vulnerability to fishing and on traits that enable the fish to reproduce before becoming captured (Cooke et al. 2007; Hollins et al. 2018; Redpath et al. 2010; Sutter et al. 2012; Uusi-Heikkilä et al. 2008).

In the context of recreational fisheries, selection is predicted to act mainly on behavior, as the vulnerability to being captured depends on fish behavior, and capture leads to either survival costs or other non-lethal fitness costs (Lennox et al. 2017; Uusi-Heikkilä et al. 2008). Vulnerability to capture by passive fishing gear, including angling, depends on traits related to risk-taking and curiosity, such as boldness and exploration tendency (Arlinghaus et al. 2017; Cooke et al. 2007; Härkönen et al. 2014; Wilson et al. 2015, reviewed in Lennox et al. (2017)), although not all studies have supported the predicted role for boldness (Louison et al. 2017; Vainikka et al. 2016). Over time, angling selection could increase the frequency of shy phenotypes in the population (Alioravainen et al. 2020a; Andersen et al. 2018; Arlinghaus et al. 2017). The shift in the behavior of fish populations may occur on top of the fishing/angling-induced decrease of body size and age-at-maturity (Bowles et al. 2020; Sharpe and Hendry 2009) and cause a personality-related decrease in resource acquisition. Eventually, these can lead to complex effects on stock recruitment (Arlinghaus et al. 2017). However, these predictions for the existence and consequences of increasing shyness due to passive fishing gear require further empirical tests.

Selection acting on personality could induce correlated metabolic effects due to physiological covariation between behaviors affecting energy balance and standard metabolic rate (SMR) (e.g., Mathot et al. 2018; Killen et al. 2011). In one of the first empirical angling selection studies, SMR was found to be 10% lower in a low-vulnerability selection line compared to a high-vulnerability selection line in largemouth bass (*Micropterus salmoides*) (Redpath et al. 2010). However, several studies have found no phenotypic association between angling vulnerability and SMR (Väätäinen et al. 2018), or between angling vulnerability and several metabolic traits, including SMR (Louison et al. 2017; Louison et al. 2018). Given that metabolic traits are are also sensitive to environmental conditions, and angling methods may impose different selection pressures in different experiments, the potential for evolutionary response in metabolic traits in response to angling-induced selection is presently not well understood.

Salmonids, just as many other taxa, can display individually distinctive behavioral strategies and coping styles (Adriaenssens and Johnsson 2011; Brelin et al. 2008; Huntingford and Adams 2005; Näslund and Johnsson 2016; Vindas et al. 2017), on which selection may act. Due to their widespread hatchery rearing, species such as the brown trout are also affected by unintended domestication, introducing shifts in life-history traits, behavior (Araki et al. 2008; Horreo et al. 2018; Huntingford 2004) and vulnerability to angling (Klefoth et al. 2013); more research on the effect of unintended domestication on fish populations used in supplemental releases is therefore warranted.

Here, our goal was to test whether angling could induce selection in behavioral traits measured under authentic predator cues or in SMR by studying one-year-old offspring of brown trout (*Salmo trutta*) from both captive and wild origins. By using replicated behavioral assays involving predator cues and collecting metabolic rate data, we add to the study by Alioravainen et al. (2020a), which focused on open-field tested personality of the offspring from the same angling experiments during their first summer. We further compared fish acclimated under 12h light: 12h dark or 24h light, because unnatural light conditions may be perceived as stressful by the fish and modify the phenotypic responses. We hypothesized that offspring from angling-vulnerable parents would have higher scores in risk-taking behavior, and higher SMR, compared to fish from non-vulnerable parents in both strains of fish.

## Material and methods

### Angling experiment and fish husbandry

> Experiments were carried out between 2015 and 2017 at the Kainuu Fisheries Research Station (www.kfrs.fi) of Natural Resources Institute Finland (Luke) under ethical license obtained from the national Animal Experiment Board in Finland (license number ESAVI/3443/04.10.07/2015). We used 1) wild, predominantly non-migratory, parental brown trout from River Vaarainjoki captured by electrofishing, and 2) captive (5–6^th^ generation), predominantly migratory brown trout from so-called Lake Oulujärvi hatchery strain. The founders of the captive brood stock came from two source populations, River Kongasjoki and River Varisjoki (Lemopoulos et al. 2019a; Alioravainen et al. 2020a). Despite originating from the same River Varisjoki watershed, only a few km apart, the captive and R. Vaarainjoki populations showed moderate genetic divergence based on fixation index (*F_ST_*-value) of 0.11 (Lemopoulos et al. 2019a). Both populations had experienced fishing pressure (mainly hook- and-line fishing) in the past, but not since R. Vaarainjoki was protected from fishing in the 1990s and the original captive population was established in the 1960s-80s.

In 2015, hatchery-origin and wild-origin adult fish were exposed to experimental fly fishing and divided into captured (high vulnerability, HV) and uncaptured (low vulnerability, LV) groups (Fig. 1) (Alioravainen et al. 2019a). Fish were fished in two size-assortative pools for each population during June and July with fly fishing gear adjusted by the size of the fish in the pools. The wild fish were fished in semi-natural 50-m^2^ ponds with a gravel-bottom outer riffle sections and *ca.* 1 m deep, concrete inner pool sections (53 and 91 visually size-sorted fish in two ponds). The hatchery fish were fished in 75-m^2^ concrete ponds with no structures (64 larger and 167 smaller fish from two different cohorts in two ponds). Angling was performed by experienced fly fishers (mainly A.V.) using unnaturally colored woolly bugger - type fly patterns tied to barbless hooks. During angling sessions, an angler fished a pond until a fish took the fly or five minutes passed, after which angling was continued at earliest one hour later. If a fish was captured, angling was continued immediately after processing, which included anesthesia with benzocaine (40 mg L^-1^), identification of passive integrated transponder (PIT) tag (Oregon RFID), or tagging when a pre-existing tag was missing, and measuring total length (to 1 mm) and weight (to 2 g). Fish that were missing PIT-tags were tagged under the skin next to the dorsal fin using 12 mm tags at this point. After processing, the fish were transferred to similar ponds (hatchery fish to a 50-m^2^ otherwise similar concrete pond) as used for each population during angling. After angling trials were finished, on 25 June 2015, all remaining wild fish that were not captured were collected by dip-netting after draining the experimental angling ponds, anaesthetized, measured and weighed (mean body lengths of fish uncaptured and captured by angling: in large fish 457 and 475 mm, respectively, and in small fish 344 and 354, respectively). Uncaptured wild fish were then combined in the same ponds as the fish captured by angling. The captured hatchery strain fish were subjected to a second round of angling ~2 weeks later to identify the most vulnerable fish, where in total eight fish were captured and prioritized for breeding the highly vulnerable line, but this was not done on wild fish due to their limited availability. Angling trials finished on 8 July 2015, and also hatchery fish were transferred back to their original ponds. Because of the warm water at the time of finishing the second round of angling, the uncaptured hatchery fish were not measured to avoid handling-induced stress and mortality. One deep-hooked small hatchery fish was found dead 5 days and one large hatchery fish 41 days after capture, but otherwise no mortality occurred between angling trials and the breeding.

**Fig. 1.**
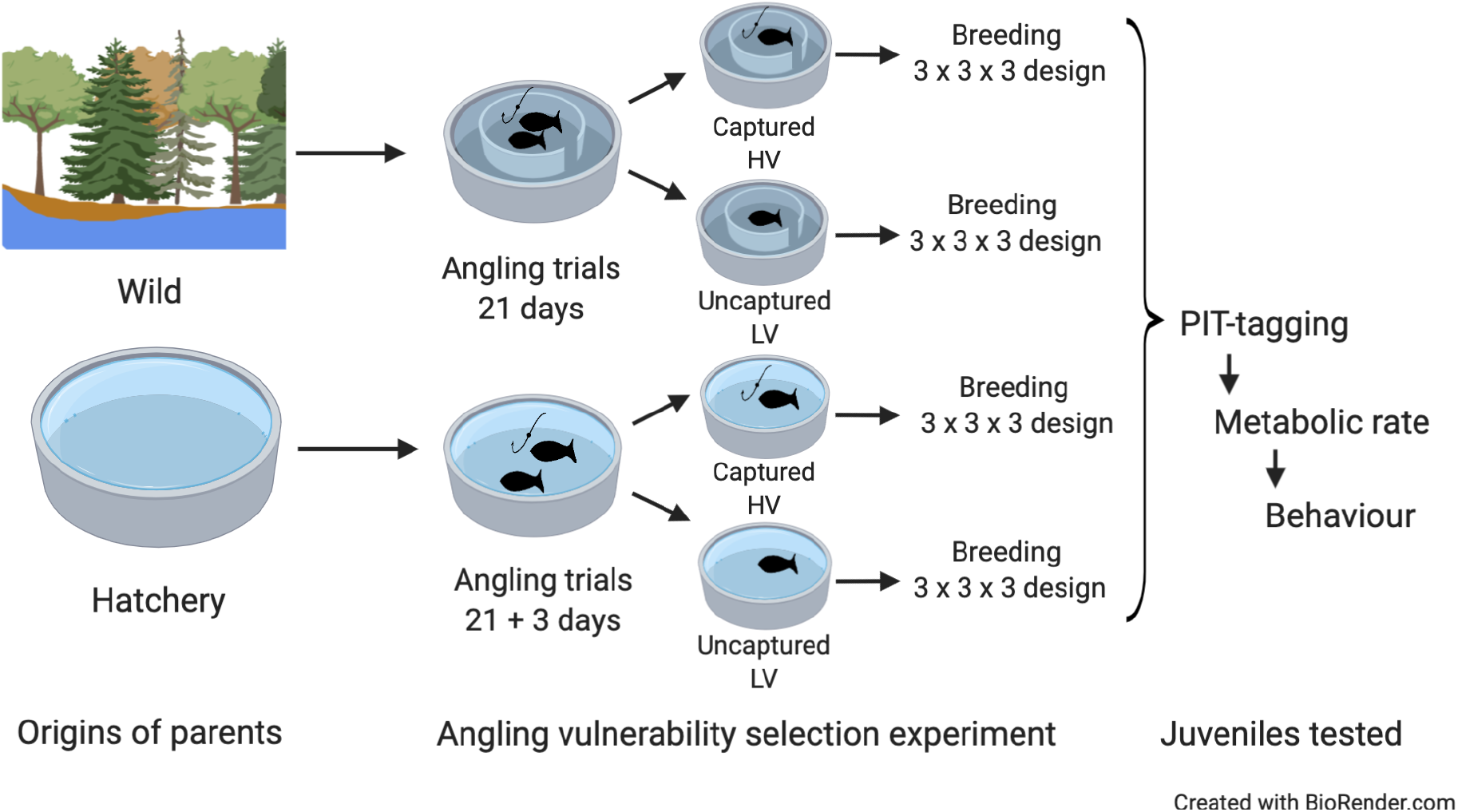
A diagram of the experimental design. Parental fish were placed in size-assortative ponds (N = 64 and 167 for large- and small-size fish in hatchery population, and N = 53 and 91 in the large- and small-size wild fish, respectively) before angling. After 21 days of angling, or after the first capture, fish were transferred to two tanks similar to those used in the angling trials. The hatchery stock individuals were thereafter fished for a second time (3 days of angling). Before breeding, fish were again combined in population-specific size-assortative ponds (not shown in diagram). Further details in Alioravainen et al. (2020a).

From the hatchery stock, 32.8 and 50.9% (corresponding to 21 and 85 fish), and from the wild stock, 28.3 and 24.2% (corresponding to 15 and 22 fish) of large and small individuals were captured, respectively. Notably, the wild fish had natural invertebrate food available in their ponds, and the structured ponds offered more hiding places, and they had clearly lower catchability than the hatchery fish (Alioravainen et al. 2020a). Very few wild fish were captured in one angling session (maximum 4) compared to the hatchery fish (maximum 11). The captured and non-captured parent fish were similar in size, indicating that vulnerability to angling was most likely related to size-independent traits (Alioravainen et al. 2020a).

The offspring used in this study were obtained from fish bred in four groups (i.e. high- and low-vulnerability [HV and LV, respectively] within each population) in the autumn of 2015. Parent fish from large and small size groups were mixed for breeding. A replicated, fully factorial 3 × 3 breeding design was used to create the F_1_-generation; males were crossed with females in all combinations in one matrix, and the matrices replicated three times for each group, details in Appendix A, and in Alioravainen et al. (2020a). In the autumn of 2016, the one-summer-old fish were tagged with individual 12-mm PIT-tags in the abdominal cavity under anesthesia (benzocaine). After tagging, the selection lines were mixed together in two 3.2 m^2^ fiberglass rearing tanks. During the whole study, fish were fed with commercial fish pellets (Raisio Oyj).

### Photoperiod acclimations

In mid-March 2017, after being reared under constant light, 100 fish were divided into two different photoperiod treatments in 0.4-m^2^ green, plastic, flow-through tanks. The tanks were covered with green nets. One treatment continued to be reared under constant light (approximately 9 lux at the water surface, n = 10/selection line/population, 40 fish in the tank), and the second treatment received a 12h light:12h dark (L:D) acclimation (approximately 12 lux during light period at the water surface, N = 15/selection line/population divided equally in two tanks, details in Appendix A).

Fish were fed using automatic belt feeders (~0.3% fish mass per day) on 5–6 days per week during approx. 4h between 8:00 and 20:00 to avoid the entrainment of endogenous rhythms by feeding. After a minimum two-week acclimation, the metabolic rate measurements were started.

### Measurement of standard metabolic rate

The SMR, i.e., post-absorptive, resting O_2_ uptake (Ṁ*O_2_*) (Nelson 2016)), of fish was measured using intermittent flow-through respirometry (Svendsen et al. 2016). The fish were not fed for 40–48h before the start of the measurement to minimize the effect of digestion on metabolic rate. As fish from the same tank were measured on multiple days, fish in each tank were fed on a rotation of 40–48h fasting followed by 1.5 d of feeding. The fish were caught by dip-netting under a dim red light into 10-L buckets, identified with a PIT-reader and transferred to the flow-through measurement chambers (diameter 33 mm, length 120 mm, Loligo Systems, Viborg, Denmark). The chambers were immersed in a water bath, which was also immersed in a flow-through tank. Measurements were started immediately and continued for approximately 23h, corresponding to 90–96 15–17-min measurement cycles for all individuals. We measured 2–4 individuals in separate horizontal glass chambers during each day of measurements. Oxygen saturation was measured in % of air saturation using two-point-calibrated DAQ-PAC-WF4 system with Sensor spot mini sensors and recorded every second in AutoResp software (Loligo Systems). Water temperature was measured using the Pt1000 temperature probe placed in the respirometer tank (Loligo Systems). Air pressure in kPa, with one decimal, was recorded daily at the start of measurements from a nearby weather station.

The respirometry measurements were started between 11:45 am and 12:10 pm by measuring the oxygen level in each empty chamber for one cycle to establish a baseline for bacterial oxygen consumption. The cycles consisted of a 6-min flush and a 9–11-min recirculation period, including a 5.5–7.5-min wait period to allow mixing of the water and a 3.5-min measurement. The temperature in the acclimation tanks was on average 3.4°C ± SD 0.12°C during the respirometer measurements, but the respirometer temperature was slightly higher than the acclimation temperature (range 3.4–4.2°C) due to the unavoidable heating of the tank by the measurements. During the measurement, the respirometer was covered with a green net, similar to what was used to cover the acclimation tanks prior to measurements, and disturbances were kept to a minimum. The fish were passive for extended periods of time during the measurements (observed from oxygen measurements). The chambers were washed using mild Deconex disinfectant and the water inside the respirometer tank was changed every 5-7 d during the measurements. Six measurements, where air bubbles were observed in the measurement chamber, were discarded from the analysis. The photoperiod was 12h:12h L:D during respirometry for all fish, because the lowest consumption was expected to occur during the dark phase.

After measurements, fish were anesthetized with benzocaine, measured for total length (to 1 mm) and weighed (to 0.1 g), after which they were transferred to new 0.4 m^2^ tanks similar to those used prior to measurements. The fish were under the same photoperiod as before the measurements and fed daily at varying times. After removing the fish, respirometer chamber oxygen levels were measured empty for one cycle to quantify bacterial respiration rates. No measurable respiration was detected without fish. The slope of the decrease in oxygen level during each 3.5-minute measurement period was calculated using linear regression with FishResp (Morozov et al. 2019) in R. An initial acclimation period was excluded by including only measurements taken after 16:00 in the analysis. We accepted all slopes where the R^2^ was > 0.9 in the calculation: this resulted in 12–70 slopes being accepted for each individual, after data from one individual with only 7 accepted slopes was excluded. R code for analyzing the raw data is provided in GitHub (see Data availability).

The SMR was calculated as the average *ṀO_2_* across all accepted cycles after comparing data obtained using several methods following Chabot et al. (2016): quantile 0.1, quantile 0.2, lowest 10 consumption values, and average. The average of accepted cycles was used as it was the only variable that showed a positive mass-dependence of metabolic rate. Other variables were not correlated with fish body mass, likely indicating inconsistencies in fish-specific accepted measurements due to variation in R^2^ values. The average of accepted cycles represents fish in post-absorptive resting conditions, excluding initial handling stress period, and thus, we expect it to be close to the real SMR of the fish.

### Setup of behavioral trials

The fish were allowed to recover from respirometry for at least four days before assayed for behavioral traits under chemical predator cues. They were not fed for 24-h prior to behavioral trials. The trials were conducted in custom-made mazes (Fig. 2) (size 400 mm wide × 1500 mm long, water depth 100 mm in the open area). During the trials, temperature in the maintenance tanks and test arenas was on average 5.0 ± SD 1.5°C. The rate of water flow was adjusted to ~8 L min^-1^ (~7.6–8.8 L min^-1^) during the trials. This allowed for at minimum 1.26 times the arena volume of water to flow between consecutive trials, which was considered sufficient to minimize potential carry-over effects of chemical cues between trials. The arena was lit by LED lights (CRI90 LED chain in waterproof silicon tube, 3000-3300K, 4.8W m^2^) situated along one long edge of the arena (>70 lux across the arena depending on the distance from the light source). Half-way across the arena was a brick gate situated next to one side, allowing entry from the other side. Behind the brick, natural pebbles (~3–5 cm in diameter) were scattered unevenly on the floor. One stone was provided for shelter and another was in the center of the arena in front of the start box (Fig. 2). Four structurally similar arenas were used in the experiment, but two of the arenas were mirror images of the other two with respect to the location of the gate.

**Fig. 2.**
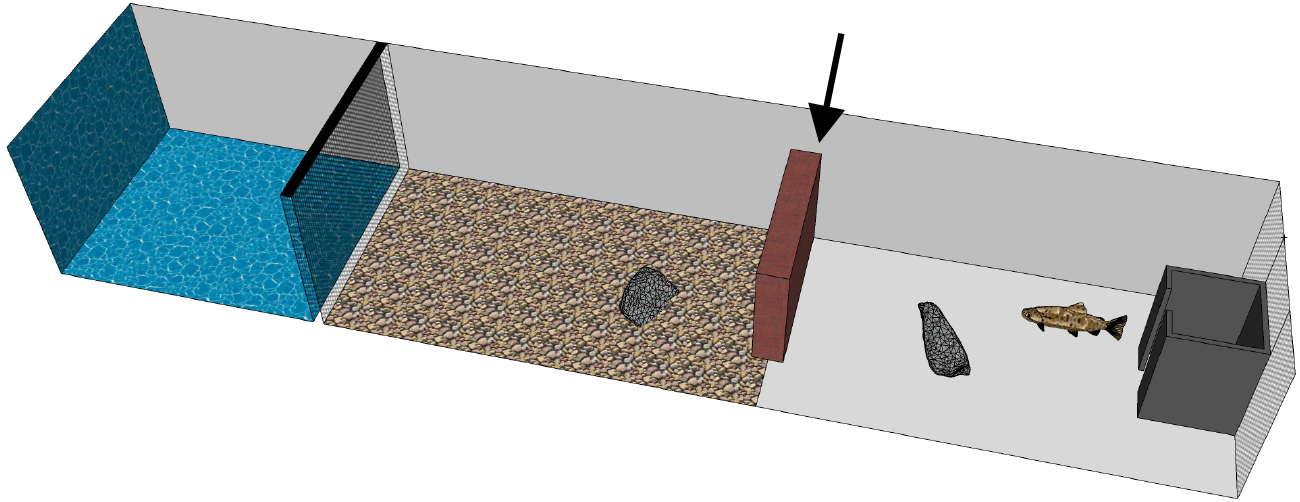
A 3D-illustration of the arena used in personality trials without the left side wall. Water flow direction is left-right. Burbot was placed in the area indicated by blue color, upstream from the net (inaccessible to the brown trout). Pebbles were scattered in the brown-colored area. The grey box indicates the start box, where the fish was placed before the start of a trial. Latency was measured as time to emerge from the box. Exploration intensity was measured as swimming activity outside the start box arena after emergence. Exploration tendency was measured as whether or not the whole body of fish passed the gate indicated by an arrow.

Upstream from the flow-through test arena section was a section divided by a metal grid (5 mm mesh size) where a hatchery-reared burbot (*Lota lota*) (length ~30-40 cm) was placed to introduce chemical cues of a natural predator of juvenile brown trout. Burbot are nocturnal bottom-dwelling predators that are likely difficult for prey to detect visually, but their odor induces antipredator responses in prey species (Ylönen et al. 2007). Burbot were maintained by feeding them with pieces of various cyprinids and vendace (*Coregonus albula*) during rearing, but fresh pieces of brown trout were used for two days prior to and every 2–3 days during the trials (feeding occurred in separate tanks, not the behavioral arenas). Approx. 5g of brown trout /meal was offered to the burbot, but not all pieces were eaten presumably due to low temperature. Burbot (N = 8) were moved to the test arenas at least one day before the trials and changed in each arena every 10–15 trials (2–3 days).

Before each trial, individual brown trout were haphazardly dip-netted from their rearing tanks under red light and placed into black 10-L buckets filled with ~8L of water from the flow-through system. Fish were identified by PIT tags and left undisturbed for 10 min before being transferred into the start box located downstream from the test arena by pouring. During each trial, the trout was acclimatized in the start box for 3 min, after which the door of the box was opened by pulling a string from behind a curtain. The movements of the fish were recorded from above using two CCTV infrared cameras (two arenas simultaneously filmed using the same camera) for 10 min (of which the first 9 min 45 s was included in the behavior analysis). The behavioral trial was repeated three times between 8:00 and 11:00 for each focal fish, with an average time of 4.3 days (range 1–8 days) between consecutive trials. One trial from four fish was omitted from the analysis due to error in data collection. The order in which batches of four fish were captured on the same day from the same tank for the four arenas was recorded (batch from hereon, levels 1–5, four individuals from batch 6/7 combined to batch 5).

### Testing behavioral responses to burbot

To confirm that burbot odor was perceived as risky by the brown trout in the personality assays, we studied the response of brown trout to burbot odor in a separate experiment using individuals from wild HV (N = 9) and wild LV groups (N = 10). These fish were acclimated in similar tanks as the personality-tested fish at 12h: 12h L:D photoperiod for one week. The behavior of each individual was tested on six different days in the presence and absence of predator odor (3 trials in each condition in haphazard order). 3–4 different arenas were used for each fish on different days to reduce fish habituation to a certain arena. All trials were conducted between 14:40 and 17:00. Control treatment arenas were emptied and thoroughly rinsed with pressurized tap water and water flow maintained for >2h before the trials to avoid carry-over effects from burbot odor in previous trials (it was not possible to conduct the tests prior to the experiment on the angling selection lines due to the limited time period with stable water temperature in the hatchery). The water used in the flow-through system originated from lake Kivesjärvi, where burbot is a common species; thus, dilute traces of burbot odor may have been present throughout the study.

### Analysis of video recordings

Behavioral data were collected from videos using manual tracking with AV Bio-Statistics 5.2 timing software. The observers were blind to the identity of fish in all recordings. Analyses were conducted in haphazard order, and each trial was analyzed once. In total four people analyzed the videos (JMP, NA, AL, SM). Four behaviors were characterized along the “exploration” personality axis, as defined by Réale et al. (2007). That is, fish were in a new environment during the tests and the odor of predator was previously unknown to the fish given the fish had not experienced natural predation risk. The behavioral events timed from the videos were 1) *latency* – the time from the start of the experiment until the whole body of fish emerged from the start box, (after Boulton et al. 2014; Moran et al. 2016; Vainikka et al. 2016); 2) time until fish passed the gate to the upstream section of the arena (arrow in Fig. 2), but this was not analyzed because of many fish not entering this section; instead we recorded 3) *exploration tendency* – a binary variable indicating whether the whole body of the fish passed the gate within the arena; and 4) *exploration intensity* – the proportion of time spent actively swimming after emerging from the start box. We used the proportion of time rather than absolute time to reduce the dependence of activity from latency. Activity was thus calculated as the total mobile time outside the box divided by total time outside the box. Fish was considered immobile when not moving for longer than ~2 s.

## Sex determination from DNA samples

To consider potential sex differences in the studied traits, we identified the sex of fish using PCR amplification of the sexually dimorphic *sdY* locus, which identifies the correct sex in brown trout with nearly 100% accuracy (Quéméré et al. 2014); details in Appendix A.

## Statistical analyses

A principal component analysis (PCA) of behavioral data collected from the fish that emerged from the box, i.e., data with no missing observations showed that eigenvalues were all < 1.5, i.e., that the three variables were not strongly correlated (Fig. S1, Fig. S2). Therefore, each behavioral variable was analyzed as separate response variable using univariate models (Table 1 for sample sizes).

**Table 1.** The number of individuals in each group in each analysis, mass-specific SMR values, and fish total body length and mass (mean ± SD) at the end of the experiment. (HV = high, LV = low vulnerability)

| Photoperiod | Population | Selection line | N (metabolic rate) | N (latency, exploration tendency & intensity) | N (body size) | N males / females (unknown) | Body length / mm | Body mass / g | 807<br>SMR mg O<sub>2</sub>/L /kg/h |
| --- | --- | --- | --- | --- | --- | --- | --- | --- | --- |
| 12:12 | Hatchery | HV | 15 | 15 | 14 | 9 / 5 (1) | 117 ± 10 | 17.1 ± 4.3 | 260.7 ± 65.5 |
| | | LV | 10 | 14 | 14 | 6 / 2 (7) | 115 ± 13 | 16.5 ± 6.7 | 300.4 ± 16.5 |
| | Wild | HV | 13 | 14 | 14 | 6 / 6 (3) | 117 ± 7 | 17.4 ± 3.2 | 261.9 ± 41.7 |
| | | LV | 15 | 15 | 14 | 7 / 7 (1) | 115 ± 9 | 16.6 ± 4.3 | 268.8 ± 49.5 |
| 24 | Hatchery | HV | 9 | 10 | 9 | 6 / 3 (1) | 119 ± 9 | 17.7 ± 4.7 | 261.2 ± 36.0 |
| | | LV | 7 | 10 | 6 | 0 / 4 (6) | 124 ± 9 | 19.6 ± 3.9 | 299.4 ± 59.4 |
| | Wild | HV | 8 | 10 | 6 | 2 / 3 (5) | 116 ± 9 | 17.8 ± 7.1 | 260.9 ± 55.3 |
| | | LV | 6 | 10 | 5 | 2 / 1 (7) | 119 ± 9 | 17.8 ± 3.0 | 305.2 ± 71.8 |

Variation in each response variable (SMR and behavioral variables) was explained with a univariate model, including breeding group and acclimation conditions as explanatory terms (Table 2). All analyses were conducted in R v.3.3.2 (R Core Team, 2016). Linear (LMM) and generalized mixed-effects models (GLMM) were fitted using package *lme4* (Bates et al. 2015) with *lmerTest* (Kuznetsova et al. 2017) and the frailty models using package *coxme* (Therneau, 2018). The data were visualized using ggplot2 (Wickham 2009). Statistical significance was determined as α = 0.05 in all models.

**Table 2.** The main statistical models used in this study. Abbreviations explained below the table.

| 810 Study section | Response variable | Model |
| --- | --- | --- |
| I. Angling selection experiment | $Log_{10}(SMR)$ | $y_{ij} = \beta_0 + \beta_1 PHO_{ij} + \beta_2 POP_{ij} + \beta_3 SEL_{ij} + \beta_4 POP_{ij} \times SEL_{ij} + \beta_5 POP_{ij} \times PHO_{ij} + \beta_7 logBM_{ij} + \beta_8 WT_{ij} + p_l + e_{ij}$ |
| | Exploration intensity | $y_{ijk} = \beta_0 + \beta_1 PHO_{ijk} + \beta_2 POP_{ijk} + \beta_3 SEL_{ijk} + \beta_4 POP_{ijk} \times SEL_{ijk} + \beta_5 POP_{ijk} \times PHO_{ijk} + \beta_6 REP_{ijk} + b_i + c_j + d_k + e_{ijk}$ |
| | Latency | $\lambda(t) = \lambda_0(t) e^{\beta_1 PHO_{ijk} + \beta_2 POP_{ijk} + \beta_3 SEL_{ijk} + \beta_4 POP_{ijk} \times SEL_{ijk} + \beta_5 POP_{ijk} \times PHO_{ijk} + \beta_6 REP_{ijk} + b_i + c_j + d_k}$ |
| | Exploration tendency (1 = explorative, 0 = un-explorative) | $y_{ij} \sim Bernoulli(p_{ijk})$<br>$logit(p_{ijk}) = \beta_0 + \beta_1 PHO_{ijk} + \beta_2 POP_{ijk} + \beta_3 SEL_{ijk} + \beta_4 POP_{ijk} \times SEL_{ijk} + \beta_5 POP_{ijk} \times PHO_{ijk} + \beta_6 REP_{ijk} + b_i + c_j + d_k$ |
| II. Behavioral responses to burbot olfactory cues | Exploration intensity | $y_{ijk} = \beta_0 + \beta_1 SEL_{ijk} + \beta_2 TRE_{ijk} + \beta_3 REP_{ijk} + \beta_4 BL_{ijk} + b_i^{(1)} CON + b_i^{(2)} BUR + c_j + d_k + e_{ijk}$ |
| | Latency | $\lambda(t) = \lambda_0(t) e^{\beta_1 SEL_{ijk} + \beta_2 TRE_{ijk} + \beta_3 REP_{ijk} + \beta_4 BL_{ijk} + b_i + c_j + d_k}$ |
| | Exploration tendency | $y_{ijk} \sim Bernoulli(p_{ijk})$<br>$logit(p_{ijk}) = \beta_0 + \beta_1 SEL_{ijk} + \beta_2 GR_{ijk} + \beta_3 REP_{ijk} + b_i + c_j + d_k$ |

$\beta_0$ Intercept, *PHO* Photoperiod, *POP* Population, *SEL* Selection, *logBM* Log<sub>10</sub> body mass in kg, *WT* Water temperature in °C, *BL* Body length in mm – mean (118.8182 mm for I, 122.4464 mm for II), *REP* Trial repeat, *TRE* Treatment, $p_l$ the random effect for chamber l, $b_i$ random effect for fish i, $c_j$ the random effect for arena j, $d_k$ the random effect for batch k, $e$ Residual, $\lambda_0$ baseline hazard, $t$ time, *CON* / *BUR* binary explanatory variables for burbot and control treatments.

SMR and body mass were log_10_-transformed for the LMM to account for the scaling of metabolic rate with body mass. The models used for different behavioral variables were: 1) LMM for *exploration intensity,* where a higher value indicates higher proportion of time spent actively swimming, 2) a frailty model (i.e. mixed effect Cox proportional hazards models for time-to event data (Collett 2015)) for *latency,* where higher trait values and negative model coefficients indicate longer time to emergence, and 3) a GLMM for *exploration tendency* (Bernoulli-distribution), where entering or not entering the upstream sector was indicated by 1 and 0, respectively. Trial repeats were encoded as a continuous variable: −1, 0, and 1 in data from angling selection experiment and as 1–6 from burbot vs. control experiment. In 8 trials, the fish jumped out of the start box prior to the trial and their behavior was analyzed for 9 min 45 s min after the jump.

For LMMs and GLMM, the main effects of population, selection line, and photoperiod were separately tested using linear hypothesis testing (function *lht* in package *car*) using restricted models, where each respective main effect and its interactions were defined zero and compared to the full model using F-tests. From LMM and GLMM models, the estimated marginal means and confidence intervals were estimated with package *ggeffects* (Lüdecke 2018). All linear models were checked for homoscedasticity and normality of residuals. For all response variables, the effect of sex was analyzed in models including the fixed effect of sex as well as the effects from original models, except photoperiod or its interactions due to limited sample size with known sex. For further details see Appendix A, and Data accessibility.

To assess individual-level correlation among the behavioral variables and SMR, Pearson’s product moment correlations were calculated between residuals from a linear model with log_10_ SMR as the response variable and log_10_ body mass in kg as the explanatory variable, and Best Linear Unbiased Predictors (BLUPs) for latency or exploration intensity. BLUPS were obtained from linear mixed models including only individual as random effect, and only for the trials where the fish emerged from the start box.

## Results

### SMR

The LMM indicated a significant effect of body mass and population on SMR, with fish from wild background having higher SMR than fish from the hatchery background, but no effect of selection line (Fig. 3, Table 3). Sex of fish had no effect on SMR (*F*_1, 58.215_ = 0.067, *P* = 0.797).

**Table 3.** Results from model for SMR. The zero levels for contrasts were: photoperiod 12:12, population hatchery, and selection line HV. F and P-values for the interactions and temperature effect were obtained from Type III sums of squares and Satterthwaite approximation for degrees of freedom. For the other fixed effects, linear hypothesis tests using F-test on restricted models with each main effect and its interactions set to zero were used – residual degrees of freedom are given for these tests. Significant (*P* < 0.05) effects shown in bold. For the intercept, t-test value is shown.

| Fixed effects | Estimate ± SE | Num<br>df | Res / Den df | F | P |
| --- | --- | --- | --- | --- | --- |
| Intercept | 0.97 ± 0.20 | 1 | 74.522 | 4.96 (t) | <0.001 |
| Photoperiod | 0.004 ± 0.02 | 2 | 72.09 | 0.41 | 0.664 |
| **Population** | **0.051 ± 0.02** | **3** | **73.355** | **2.78** | **0.047** |
| Selection | 0.031 ± 0.02 | 2 | 72.281 | 1.27 | 0.286 |
| Temperature | 0.02 ± 0.02 | 1 | 73.866 | 1.32 | 0.254 |
| **Log<sub>10</sub> body mass** | **0.27 ± 0.09** | **1** | **72.773** | **9.90** | **0.0024** |
| Pop × selection | -0.031 ± 0.03 | 1 | 72.173 | 1.342 | 0.250 |
| Pop × photoperiod | 0.014 ± 0.03 | 1 | 72.119 | 0.241 | 0.625 |
| **Random effects** | **Variance (SD<sup>2</sup>)** |  |  |  |  |
| Chamber | 0.039<sup>2</sup> |  |  |  |  |
| Residual | 0.060<sup>2</sup> |  |  |  |  |

**Fig. 3.**
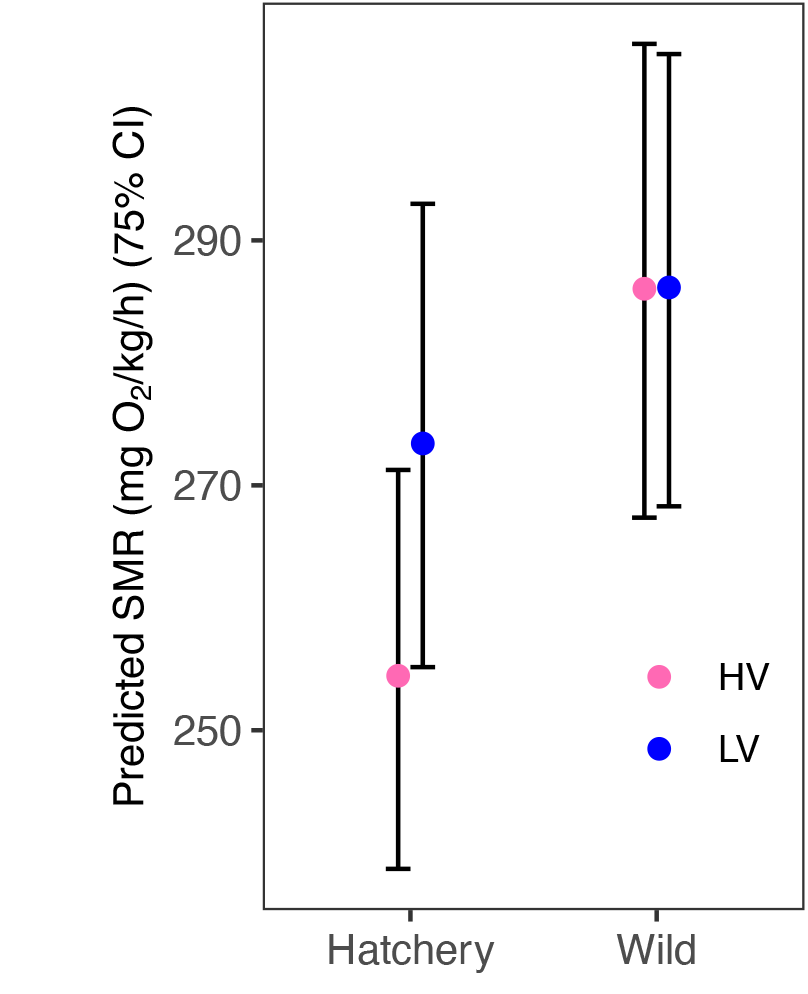
Predicted SMR (estimated marginal means) across two populations and angling selection lines (HV – high vulnerability, LV - low vulnerability) in brown trout. Data from two photoperiods combined. Estimates and 75% confidence intervals were obtained from a linear mixed model and conditioned on fish average body mass and average temperature during measurements, back-transformed to linear scale and divided by average body size in kg to obtain mg O_2_kg^-1^h^-1^. For N in each group, see Table 1.

### Behavior in angling selection lines

Fish emerged from the start box during the recorded time in ~84% of the trials. Mean latency for the fish that emerged was 1.83 min (range 0 – 9.54 min). There was a significant effect of angling selection (*P* = 0.027) and a slightly non-significant interaction effect (*P* = 0.054) of population background and angling selection on latency (Table 4). This was observed as an elevated risk to emerge in fish from LV background compared to HV background in the hatchery population, but not in the wild population (Fig. 4A).

**Table 4.**
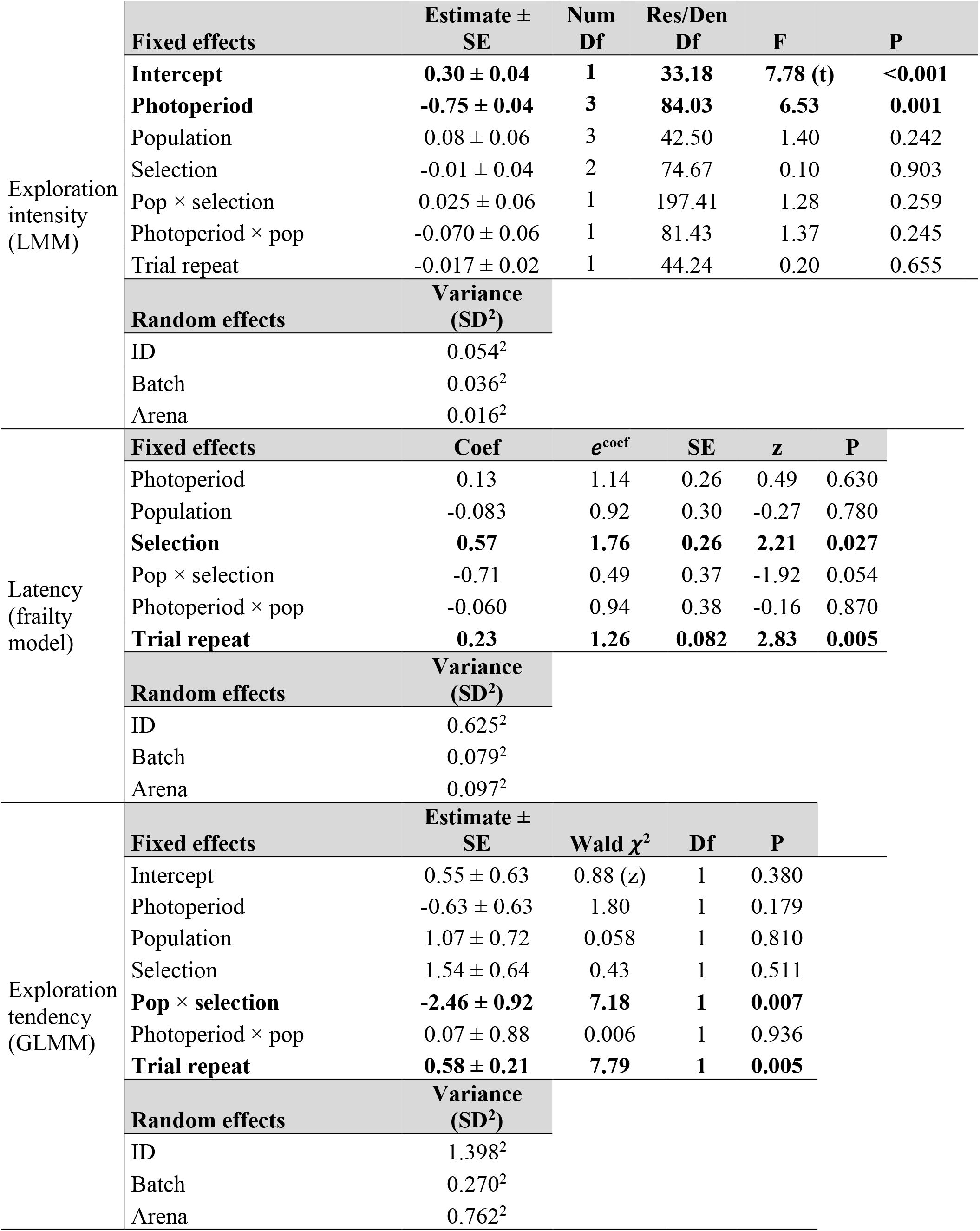
Results of models for behavior traits in brown trout from hatchery and wild populations and two angling selection lines (HV and LV). The zero levels for contrasts in all models were: photoperiod 12:12, population hatchery, and selection line HV. For model equations, see Table 2. For exploration intensity, F- and P-values for the interactions and trial repeat were obtained from Type III test, and for the other main effects from linear hypothesis tests using restricted models with each main effect and its interactions set to zero. For latency, proportional hazard estimates for risk or emergence (± standard error) are shown with hazard ratios (*e^coef^*). For latency and exploration tendency, Wald Chisquare test was used to determine significance of fixed effects. Fixed effects with *P* < 0.05 shown in bold. For intercepts, t- or z-test values shown.

**Fig. 4.**
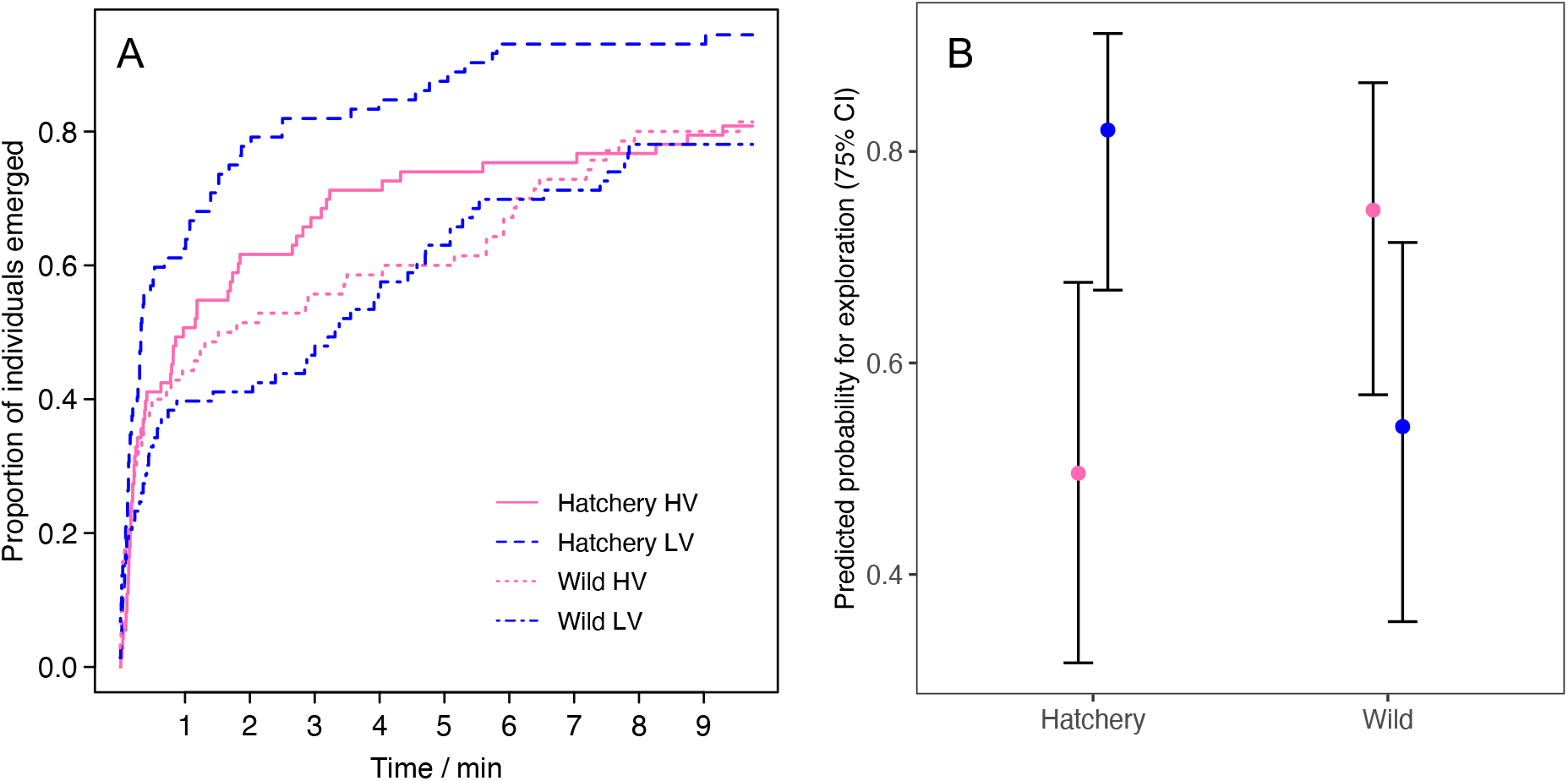
Behavioral differences between two angling vulnerability selection lines (HV – high vulnerability, LV – low vulnerability) within the hatchery and wild populations. Data from photoperiods combined for clarity. A) Curves showing the proportion of individuals emerged from the start box, drawn with Kaplan-Meier estimator. B) Predicted exploration tendency (estimated marginal means) from GLMM with 75% confidence intervals for predicted values. Angling selection had opposing effects on exploration tendency in the two populations (Table 4). Predictions were made for the first trial repeat. For N in each group, see Table 1. Pink = HV, blue = LV.

Exploration intensity did not differ between populations or angling selection lines, but the fish spent less time exploring the arena after acclimation in constant light compared to the 12h light:12h dark photoperiod (Table 4). Mean exploration intensity was 0.37 (range 0.0015 – 1.0).

Angling selection had contrasting effects on exploration tendency in each population: in the hatchery population, a higher proportion of fish from LV selection line were explorative than from HV selection line, while there was an opposite tendency in the wild population (Fig. 4B; Table 4). In addition, exploration tendency increased with repeats of the behavioral trial.

Sex did not have a significant effect on any behavioral trait (female vs. male, Exploration intensity: *F*_1,39.612_ = 1.217, *P* = 0.277; Latency: *e^coef^* = 1.03, *z* = 0.29, *P* = 0.770; Exploration tendency: *z* = −0.514, *P* = 0.607).

There was no correlation between mass-adjusted SMR and behavioral traits as assessed with initial, anti-conservative test using individual BLUPs (see Houslay & Wilson 2017) for neither exploration intensity (r = −0.006, t = −0.0579, P = 0.96) nor latency (r = −0.083, t = −0.75_79_, P = 0.46).

### Behavioral responses to predator presence

The were no significant direct effects of predator cues on behavioral traits (Table 5). Repeating the behavior trials six times for each fish, three times with a burbot present and three times in control conditions, led to a significant decrease in exploration intensity with trial repeats (Table 5). The variance of exploration intensity between individuals appeared seemingly higher in the presence of burbot, but this was not significant in Levene’s test of homogeneity of variance (*F*_1,93_ = 0.214, *P* = 0.645). Risk of emergence increased with increasing behavior trial repeats and between-individual variation in latency was high (~10% higher variance in burbot vs. control data compared to data from angling selection lines).

**Table 5.**
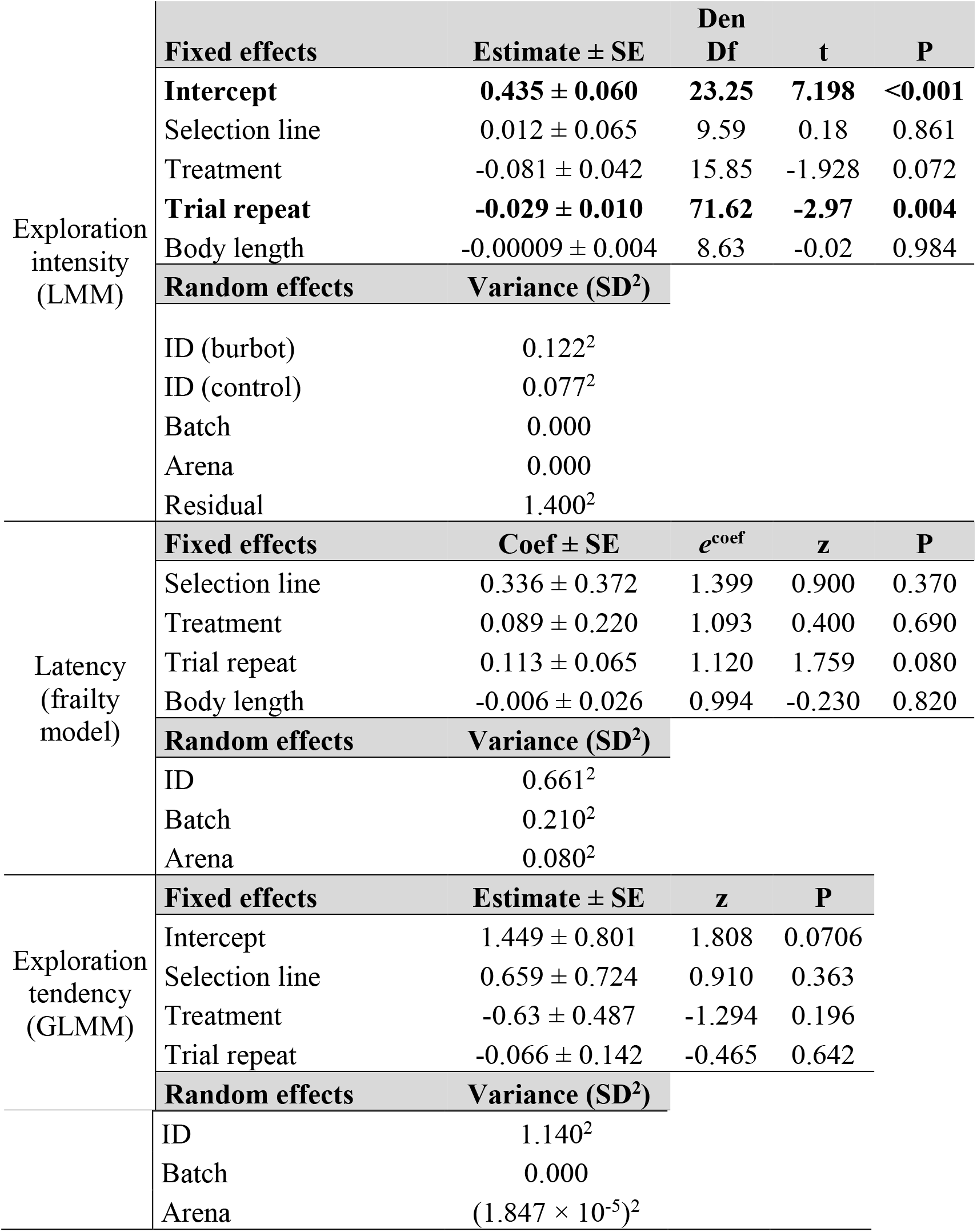
Results of models for behavioral traits in the presence of predatory olfactory cues and control conditions in brown trout. For exploration intensity, the t-test was used with Satterthwaite approximations to degrees of freedom and the model was fit with restricted maximum likelihood. For latency, proportional hazard estimates for risk or emergence (± standard error) are shown with hazard ratios (*e^coef^*). For latency and exploration tendency, Wald Chisquare test was used to determine significance of fixed effects. The zero levels for contrasts in all models were: treatment control and selection line HV. Significant effects (*P* < 0.05) shown in bold.

| | Fixed effects | Estimate ± SE | Den<br>Df | t | P |
| --- | --- | --- | --- | --- | --- |
| Exploration<br>intensity<br>(LMM) | **Intercept** | **0.435 ± 0.060** | **23.25** | **7.198** | **<0.001** |
| | Selection line | 0.012 ± 0.065 | 9.59 | 0.18 | 0.861 |
| | Treatment | -0.081 ± 0.042 | 15.85 | -1.928 | 0.072 |
|  | **Trial repeat** | **-0.029 ± 0.010** | **71.62** | **-2.97** | **0.004** |
| | Body length | -0.00009 ± 0.004 | 8.63 | -0.02 | 0.984 |
|  | **Random effects** | **Variance (SD<sup>2</sup>)** |  |  |  |
|  | ID (burbot) | 0.122<sup>2</sup> |  |  |  |
|  | ID (control) | 0.077<sup>2</sup> |  |  |  |
|  | Batch | 0.000 |  |  |  |
|  | Arena | 0.000 |  |  |  |
|  | Residual | 1.400<sup>2</sup> |  |  |  |
| Latency<br>(frailty<br>model) | **Fixed effects** | **Coef ± SE** | **$e^{coef}$** | **z** | **P** |
| | Selection line | 0.336 ± 0.372 | 1.399 | 0.900 | 0.370 |
| | Treatment | 0.089 ± 0.220 | 1.093 | 0.400 | 0.690 |
| | Trial repeat | 0.113 ± 0.065 | 1.120 | 1.759 | 0.080 |
| | Body length | -0.006 ± 0.026 | 0.994 | -0.230 | 0.820 |
|  | **Random effects** | **Variance (SD<sup>2</sup>)** |  |  |  |
|  | ID | 0.661<sup>2</sup> |  |  |  |
|  | Batch | 0.210<sup>2</sup> |  |  |  |
|  | Arena | 0.080<sup>2</sup> |  |  |  |
| Exploration<br>tendency<br>(GLMM) | **Fixed effects** | **Estimate ± SE** | **z** | **P** |  |
| | Intercept | 1.449 ± 0.801 | 1.808 | 0.0706 | |
| | Selection line | 0.659 ± 0.724 | 0.910 | 0.363 | |
| | Treatment | -0.63 ± 0.487 | -1.294 | 0.196 | |
| | Trial repeat | -0.066 ± 0.142 | -0.465 | 0.642 | |
|  | **Random effects** | **Variance (SD<sup>2</sup>)** |  |  |  |

|  |  |
| --- | --- |
| ID | $1.140^2$ |
| Batch | 0.000 |
| Arena | $(1.847 \times 10^{-5})^2$ |

## Discussion

We found that captured and non-captured brown trout produced offspring that differed in exploration-related behaviors. Against our expectations, the LV selection line was more explorative in a new environment than the HV line in the hatchery population. However, in the wild population, there was no significant response to angling selection in exploration measured as latency to emerge. However, we found a significantly contrasting response in the likelihood of exploring the arena after emergence compared to the hatchery line, with the HV line being more explorative than the LV line. The results broadly agree with behavioral responses found in the offspring from the same angling selection experiment in their first summer, although the performed personality assays differed between the two studies;

Alioravainen et al. (2020a) assayed fingerlings at warm temperatures in simple arenas without predator cues. Together, these studies provide evidence for a heritable link between angling vulnerability and exploratory behavior assessed in controlled settings but highlight that such connections can be population or environment specific. The smaller differences in the behavior between angling selection lines from the wild population than from the hatchery population may result from several reasons. First, genetic variation in the wild population was smaller than in the hatchery population (Lemopoulos et al. 2019a). Second, the parental wild fish had no juvenile hatchery history, and they were captured directly from seminatural ponds with a stream and a pool section during angling, while the hatchery fish were fished in a plain concrete tank under high density, which may have introduced density-dependent behavioral effects that were absent in the wild fish.

The result that mass-corrected SMR was higher in the offspring of wild fish than those from hatchery parents was unexpected. The hatchery environment is expected to favor individuals with a fast metabolic rate (Reid et al. 2012; Robertsen et al. 2018). However, because metabolic rate measurements can induce stress responses (Murray et al. 2017) and domestication-related phenotypic changes can be evident in hatchery environments already after a single generation in salmonids (Christie et al. 2016, Islam et al. 2020), the result could be explained by a better ability of the hatchery-population fish to adjust to handling during metabolic rate measurements. Further, metabolic rate might relate to the life-history strategy of the fish (Rosenfeld et al. 2015), as the wild fish represented resident and the captive fish migratory brown trout forms (Lemopoulos et al. 2019b). The potential effects of angling selection on metabolic rate were smaller than the effects of population, and our technique lacked sensitivity to detect minor differences between groups, but this could be reassessed in further studies.

Differences in stress coping styles, i.e. sensitivity of the neuroendocrine stress responses (Schjolden et al. 2005; Koolhaas et al. 2010), could partly explain why the results on behaviors and SMR were partly contradicting our expectations. If the LV fish from the hatchery population were more reactive compared to the HV fish, their behavior may have indicated a higher stress response to the experiment and heightened escape behavior; this effect has also been suggested to occur in pike (*Esox lucius*) (Laskowski et al. 2016). Angling intensity itself can also influence the serotonergic and dopamine systems in fish (Koeck et al. 2018), and given that Koeck et al. (2018) found clear species differences in the effect between rainbow trout and brown trout, it can be speculated that population differences in how fish respond physiologically to disturbance caused by angling are also possible. Changes in stress-related physiological responses could explain which individuals are most vulnerable to angling. Relatedly, the most angling-vulnerable parent fish in the hatchery strain may have had the lowest status in the dominance hierarchy within the concrete ponds, and therefore been the hungriest and the most prone to attack the fly patterns. In contrast to the hatchery population, wild fish were under more natural-like ponds during the angling trials and this may have allowed to them to maintain more of their natural behavior, including hierarchy related to feeding, and by extension, to attacking the flies.

It is likely that for both population and angling selection line differences that genetic inheritance would explain our results at least partly; Ågren et al. (2019) showed that the heritability in exploration-related principal component was 0.1 ± 0.065 SE in the same hacthery strain of brown trout, along with hatchery-wild crossbred strains. In Kortet et al. (2014), low heritabilities were reported for exploration and boldness-related personality axis, but that of freezing tendency was 0.14, similar to Ågren et al (2019). In three-spined stickleback (*Gasterosteus aculeatus*), behavioral traits related to exploration and boldness in the presence and absence of predation risk were moderately heritable (Dingemanse et al. 2009). Importantly, angling vulnerability itself is heritable in largemouth bass (Philipp et al. 2009).

Photoperiod had no influence on the relative differences between populations or on the absolute metabolic and behavioral traits, except on exploration intensity, which, however, was not affected by population background or angling selection. This suggests that exploration and metabolic rate differences between populations can be consistent across environments. Constant light is not encountered by brown trout during the winter; hence the 24-hour light regime could be considered unnatural and potentially stressful for the fish. Constant light can disrupt entrainment of endogenous rhythms by inhibiting the synthesis of melatonin and by directly affecting photosensitive proteins (Falcón et al. 2010; Peirson et al. 2009). In general, non-tropical species are expected to be particularly sensitive to photoperiod disturbances due to the role of day length in anticipating seasonal changes in environmental conditions (Borniger et al. 2017).

The differences we found between populations can also be explained by the level of domestication, as the hatchery stock had been reared in captivity for several generations. Populations frequently differ in e.g., metabolic rate and behavioral syndromes (Dingemanse et al. 2007; Lahti et al. 2002; Polverino et al. 2018), driven by environmental differences, natural selection, founder effects, and genetic drift. The populations we studied also differed in their life-histories, with the wild population being clearly less migratory than the hatchery population (Lemopoulos et al. 2019b). Moreover, juveniles from the wild population show lower tendency for post-release dispersal in a stream environment than juveniles from the hatchery population (Alioravainen et al. 2020b). Although we reared offspring under common garden conditions and maximized genetic diversity within each group through a fully factorial breeding matrix, it is possible that differences in the early rearing environments of wild and hatchery parents had contrasting effects on offspring through parental or epigenetic effects (Crews et al. 2012; Reddon 2012). The duration of these effects on offspring physiology can be short- or long-lasting (Bell et al. 2016; Metzger and Schulte 2016; Munday et al. 2017).

Our goal was to study exploration/risk-taking-related behaviors by subjecting fish to the olfactory cues of a natural predator that had fed on conspecifics, which was expected to cause a strong antipredator response (Vilhunen and Hirvonen 2003; Ferrari et al. 2010). Although no direct effect of predator presence on brown trout behavior was found in this study, we found a strong decline in the exploration intensity of fish with increasing predator/control trial repeats, which could indicate the development of an antipredator response, evident as increased hiding and decreased activity in juvenile salmonids (Vilhunen 2006; Vilhunen and Hirvonen 2003; Kopack et al. 2015). However, we did not separate fish into control and predator exposure treatments, but each fish in the test served as their own control, and thus the effect may also be related to habituation into the test arenas. We conducted the test of predator cues only on the wild fish offspring, given that innate responses were expected to be stronger in this population, but notably none of the individuals in the behavior trials in this study had been exposed to predators before the trials, apart from potential traces of piscivore odors in the rearing water. It is therefore not surprising that results were not as strong as in previous studies using wild-caught individuals (Álvarez and Nicieza 2003). In our study, the scarcity of responses to the presence of predator odor, measured in the offspring of wild fish, indicate only weak innate responses. Nevertheless, the (non-significant) tendency for lower exploration intensity in the presence of burbot than in control conditions resembles previously shown antipredator responses in salmonids.

## Conclusions

Our results suggest population-specific potential for rapid human-induced evolution in the behavior of a popular fishing target species. Population differences in the response to selection may have arisen from contrasting dependence of angling vulnerability from fish behavior, or from differences in the heritability of selected behaviors. This highlights the complexity of ecological and innate factors that can contribute to angling-induced selection in natural populations. Overall, the results from this common garden experiment suggest a significant genetic effect upon the behavior of brown trout parr.

## Data accessibility

All data and R codes for analyzing respirometry data and for the statistical models are available in GitHub (https://github.com/jprokkola/Strutta_repo). Data will additionally be made available in a permanent repository upon acceptance for publication. Videos of behavior trials will be made publicly available in Figshare (accession) upon acceptance for publication.

## Appendices

## Appendix A. Supplemental material and methods

## Author contributions

A.V. and P.H. produced the selection lines, J.M.P, N.A. and A.V. designed the experiment, J.M.P., N.A., S.M. and A.L. collected the data, J.M.P. and L.M. analyzed the data, J.M.P. wrote the initial draft of the manuscript. All authors contributed to preparing the manuscript.

## Acknowledgements

We thank the staff of Kainuu Fisheries Research Station for their help in catching, breeding and rearing fish, Dr. Hannu Huuskonen for advice in setting up the respirometry, Dr. Chris Elvidge for comments on the manuscript, and Prof. Robert Arlinghaus for participation in the planning of the selection lines. J.M.P., A.V. and A.L. were supported by the Academy of Finland grant for A.V. (nr. 286261). J.M.P. was also supported by Oskar Öflund’s foundation and by the Finnish Cultural Foundation.

## Literature Cited

Adriaenssens B, Johnsson JI. 2011. Shy trout grow faster: exploring links between personality and fitness-related traits in the wild. Behav Ecol 22:135–143. doi: 10.1093/beheco/arq185

Alioravainen N, Hyvärinen P, Vainikka A. 2020a. Behavioural effects in juvenile brown trout in response to parental angling selection. Can J Fish Aquat Sci 77: 365–374, https://doi.org/10.1139/cjfas-2018-0424

Alioravainen N, Prokkola JM, Lemopoulos A, Härkönen L, Hyvärinen P, Vainikka A. 2020b. Postrelease exploration and diel activity of hatchery, wild, and hybrid strain brown trout in seminatural streams. Can J Fish Aquat Sci 0:1–8. https://doi.org/10.1139/cjfas-2019-0436

Allendorf FW, Hard JJ. 2009. Human-induced evolution caused by unnatural selection through harvest of wild animals. Proc Natl Acad Sci USA 106:9987–9994. doi: 10.1073/pnas.0901069106

Álvarez D, Nicieza AG. 2003. Predator avoidance behaviour in wild and hatchery-reared brown trout: the role of experience and domestication. J Fish Biol 63:1565–1577. doi: 10.1111/j.1095-8649.2003.00267.x

Andersen KH, Marty L, Arlinghaus R. 2018. Evolution of boldness and life history in response to selective harvesting. Can J Fish Aquat Sci 75:271–281. doi: 10.1139/cjfas-2016-0350

Araki H, Berejikian BA, Ford MJ, Blouin MS. 2008. Fitness of hatchery-reared salmonids in the wild. Evol Appl 1:342–355. doi: 10.1111/j.1752-4571.2008.00026.x

Arlinghaus R, Laskowski KL, Alós J, Klefoth T, Monk CT, Nakayama S, Schröder A. 2017. Passive gear-induced timidity syndrome in wild fish populations and its potential ecological and managerial implications. Fish Fish 18:360–373. doi: 10.1111/faf.12176

Bates D, Maechler M, Bolker B, Walker S. 2015. Fitting Linear Mixed-Effects Models Using lme4. J Stat Softw, 67:1–48. doi:10.18637/jss.v067.i01

Bell AM, Laura KEM, Stein LR. 2016. Effects of mothers’ and fathers’ experience with predation risk on the behavioral development of their offspring in threespined sticklebacks. Curr Opin Behav Sci 7:28–32. doi: 10.1016/j.cobeha.2015.10.011

Borniger JC, Cisse YM, Nelson RJ, Martin LB. 2017. Seasonal Variation in Stress Responses. Pp 411–419 in Fink, G., ed. Stress: Neuroendocrinology and Neurobiology, Amsterdam: Elsevier/AP. doi: 10.1016/b978-0-12-802175-0.00041-3

Boulton K, Grimmer AJ, Rosenthal GG, Walling CA, Wilson AJ. 2014. How stable are personalities? A multivariate view of behavioural variation over long and short timescales in the sheepshead swordtail, *Xiphophorus birchmanni*. Behav Ecol Sociobiol 68:791–803. doi: 10.1007/s00265-014-1692-0

Bowles, E, Marin, K, Mogensen, S, MacLeod, P, Fraser, DJ. 2020. Size reductions and genomic changes within two generations in wild walleye populations: associated with harvest?. Evol Appl 13: 1128–1144. https://doi.org/10.1111/eva.12987

Brelin D, Petersson E, Dannewitz J, Dahl J, Winberg S. 2008. Frequency distribution of coping strategies in four populations of brown trout *(Salmo trutta)*. Horm Behav 53:546–556. doi: 10.1016/j.yhbeh.2007.12.011

Chabot, D., Steffensen, J.F. and Farrell, A.P. 2016. The determination of standard metabolic rate in fishes. J Fish Biol, 88: 81–121. https://doi.org/10.1111/jfb.12845

Christie MR, Marine ML, Fox SE, French RA, Blouin MS. 2016. A single generation of domestication heritably alters the expression of hundreds of genes. Nat Commun. 7:10676. doi:10.1038/ncomms10676

Collett D. 2015. Modelling Survival Data in Medical Research, Third Edition. Chapman & Hall/CRC Texts in Statistical Science.

Coltman DW, O’Donoghue P, Jorgenson JT, Hogg JT, Strobeck C, Festa-Bianchet M. 2003. Undesirable evolutionary consequences of trophy hunting. Nature 426:655–658. doi: 10.1038/nature02177

Cooke SJ, Suski CD, Ostrand KG, Wahl DH, Philipp DP. 2007. Physiological and behavioral consequences of long-term artificial selection for vulnerability to recreational angling in a teleost fish. Physiol Biochem Zool 80:480–490. doi: 10.1086/520618

Crews D, Gillette R, Scarpino SV, Manikkam M, Savenkova MI, Skinner MK. 2012. Epigenetic transgenerational inheritance of altered stress responses. Proc Natl Acad Sci USA 109:9143–9148. doi: 10.1073/pnas.1118514109

Dingemanse NJ, Van der Plas F, Wright J, Réale D, Schrama M, Roff DA, Van der Zee E, Barber I. 2009. Individual experience and evolutionary history of predation affect expression of heritable variation in fish personality and morphology. Proc Biol Sci 276:1285–1293. doi: 10.1098/rspb.2008.1555

Dingemanse NJ, Wright J, Kazem AJN, Thomas DK, Hickling R, Dawnay N. 2007. Behavioural syndromes differ predictably between 12 populations of three-spined stickleback. J Anim Ecol 76:1128–1138. doi: 10.1111/j.1365-2656.2007.01284.x

Falcón J, Migaud H, Muñoz-Cueto JA, Carrillo M. 2010. Current knowledge on the melatonin system in teleost fish. Gen Comp Endocrinol 165:469–482. doi: 10.1016/j.ygcen.2009.04.026

Fugère V, Hendry AP. 2018. Human influences on the strength of phenotypic selection. Proc Natl Acad Sci USA 115:10070–10075. doi: 10.1073/pnas.1806013115

Härkönen L, Hyvärinen P, Paappanen J, Vainikka A. 2014. Explorative behavior increases vulnerability to angling in hatchery-reared brown trout. *Salmo trutta)*. Can J Fish Aquat Sci 71:1900–1909. doi: 10.1139/cjfas-2014-0221

Hollins J, Thambithurai D, Koeck B, Crespel A, Bailey DM, Cooke SJ, Lindström J, Parsons KJ, Killen SS. 2018. A physiological perspective on fisheries-induced evolution. Evol Appl 11:561–576. doi: 10.1111/eva.12597

Horreo JL, Valiente AG, Ardura A, Blanco A, Garcia-Gonzalez C, Garcia-Vazquez E. 2018. Nature versus nurture? Consequences of short captivity in early stages. Ecol Evol 8:521–529. doi: 10.1002/ece3.3555.

Houslay TM, Wilson AJ. 2017. Avoiding the misuse of BLUP in behavioural ecology, Behav Ecol 28, 948–952, doi: https://doi.org/10.1093/beheco/arx023

Huntingford F, Adams C. 2005. Behavioural syndromes in farmed fish: implications for production and welfare. Behaviour 142:1207–1221.

Huntingford FA. 2004. Implications of domestication and rearing conditions for the behaviour of cultivated fishes. J Fish Biol 65:122–142. doi: 10.1111/j.1095-8649.2004.00562.x

Islam SS, Yates MC, Fraser DJ. 2020. Single generation exposure to a captive diet: a primer for domestication selection in a salmonid fish? bioRxiv 2020.01.24.919175; doi: https://doi.org/10.1101/2020.01.24.919175

Johnsson JI, Näslund J. 2018. Studying behavioural variation in salmonids from an ecological perspective: observations questions methodological considerations. Rev Fish Biol Fish 28:795–823. doi: 10.1007/s11160-018-9532-3

Kern EMA, Robinson D, Gass E, Godwin J, Langerhans RB. 2016. Correlated evolution of personality, morphology and performance. Anim Behav 117:79–86. doi: 10.1016/j.anbehav.2016.04.007

Killen SS, Marras S, McKenzie DJ. 2011. Fuel, fasting, fear: routine metabolic rate and food deprivation exert synergistic effects on risk-taking in individual juvenile European sea bass. J Anim Ecol 80:1024–1033. doi: 10.1111/j.1365-2656.2011.01844.x

Klefoth T, Pieterek T, Arlinghaus R. 2013. Impacts of domestication on angling vulnerability of common carp, *Cyprinus carpio:* the role of learning, foraging behaviour and food preferences. Fisheries Management and Ecology 20:174–186. doi: 10.1111/j.1365-2400.2012.00865.x

Koeck B, Závorka L, Aldvén D, Näslund J, Arlinghaus R, Thörnqvist P-O, Winberg S, Björnsson BT, Johnsson J. 2018. Angling selects against active and stress-resilient phenotypes in rainbow trout. Can J Fish Aquat Sci. 76. 10.1139/cjfas-2018-0085.

Koolhaas JM, de Boer SF, Coppens CM, Buwalda B. 2010. Neuroendocrinology of coping styles: Towards understanding the biology of individual variation. Front Neuroendocrinol 31:307–321. doi: 10.1016/j.yfrne.2010.04.001

Kopack CJ, Broder ED, Lepak JM, Fetherman ER, Angeloni LM. 2015. Behavioral responses of a highly domesticated, predator naive rainbow trout to chemical cues of predation. Fish Res 169:1–7. doi: 10.1016/j.fishres.2015.04.005

Kortet R, Vainikka A, Janhunen M, Piironen J, Hyvärinen P. 2014. Behavioral variation shows heritability in juvenile brown trout *Salmo trutta*. Behav Ecol Sociobiol 68: 927–934. https://doi.org/10.1007/s00265-014-1705-z

Kuznetsova A, Brockhoff PB, Christensen RHB. 2017). lmerTest Package: Tests in Linear Mixed Effects Models. J Stat Softw, 82:1–26. doi: 10.18637/jss.v082.i13.

Lahti K, Huuskonen H, Laurila A, Piironen J. 2002. Metabolic rate and aggressiveness between Brown Trout populations. Funct Ecol 16:167–174. doi: 10.1046/j.1365-2435.2002.00618.x

Laskowski KL, Monk CT, Polverino G, Alos J, Nakayama S, Staaks G, Mehner T, Arlinghaus R. 2016. Behaviour in a standardized assay, but not metabolic or growth rate, predicts behavioural variation in an adult aquatic top predator *Esox lucius* in the wild. J Fish Biol 88:1544–1563. doi: 10.1111/jfb.12933

Lemopoulos A, Prokkola JM, Uusi-Heikkilä S, Vasemägi A, Huusko A, Hyvärinen P, Koljonen ML, Koskiniemi J, Vainikka A. 2019a. Comparing RADseq and microsatellites for estimating genetic diversity and relatedness Implications for brown trout conservation. Ecol Evol 9:2106–2120. doi: 10.1002/ece3.4905

Lemopoulos A, Uusi-Heikkilä S, Hyvärinen P, Alioravainen N, Prokkola JM., Elvidge C, Vasemägi A, Vainikka A. 2019b. Association mapping based on a common-garden migration experiment reveals candidate genes for migration tendency in brown trout. G3: Genes, Genom, Genet 9:2887–2896. doi: https://doi.org/10.1534/g3.119.400369

Lennox RJ, Alos J, Arlinghaus R, Horodysky A, Klefoth T, Monk CT, Cooke SJ. 2017. What makes fish vulnerable to capture by hooks? A conceptual framework and a review of key determinants. Fish Fish 18:986–1010. doi: 10.1111/faf.12219

Louison MJ, Adhikari S, Stein JA, Suski CD. 2017. Hormonal responsiveness to stress is negatively associated with vulnerability to angling capture in fish. J Exp Biol 220:2529–2535. doi: 10.1242/jeb.150730

Louison MJ, Stein JA, Suski CD. 2018. Metabolic phenotype is not associated with vulnerability angling in bluegill sunfish *(Lepomis macrochirus)*. Can J Zool 96:1264–1271. doi: 10.1139/cjz-2017-0363

Lüdecke D. 2018. ggeffects: Tidy Data Frames of Marginal Effects from Regression Models. Journal of Open Source Software 3: 772. doi: 10.21105/joss.00772

Mathot KJ, Dingemanse NJ, Nakagawa S. 2018. The covariance between metabolic rate and behaviour varies across behaviours and thermal types: meta-analytic insights. Biol Rev 4:1056–1074. doi:10.1111/brv.12491

Ferrari M.C.O., Wisenden B.D., Chivers D.P. 2010. Chemical ecology of predator–prey interactions in aquatic ecosystems: a review and prospectus. Can J Zool. 88: 698–724. https://doi.org/10.1139/Z10-029

Metzger DCH, Schulte PM. 2016. Maternal stress has divergent effects on gene expression patterns in the brains of male and female threespine stickleback. Proc Biol Sci 283. doi: 10.1098/rspb.2016.1734

Moran NP, Mossop KD, Thompson RM, Wong BBM. 2016. Boldness in extreme environments: temperament divergence in a desert-dwelling fish. Anim Behav 122:125–133. doi: 10.1016/j.anbehav.2016.09.024

Morozov, S., McCairns, R.J.S., Merila, J. 2019 FishResp: R package and GUI application for analysis of aquatic respirometry data. Conserv Physiol 7: coz003; https://doi.org/10.1093/conphys/coz003

Munday PL, Donelson JM, Domingos JA. 2017. Potential for adaptation to climate change in a coral reef fish. Global Chang Biol 23:307–317. doi: 10.1111/gcb.13419

Murray L, Rennie MD, Svendsen JC, Enders EC. 2017. Respirometry increases cortisol levels in rainbow trout *Oncorhynchus mykiss:* implications for measurements of metabolic rate. J Fish Biol 90: 2206–2213. doi:10.1111/jfb.13292

Näslund J, Johnsson JI. 2016. State-dependent behavior and alternative behavioral strategies in brown trout *(Salmo trutta* L.) fry. Behav Ecol Sociobiol 70:2111–2125. doi: 10.1007/s00265-016-2215-y

Nelson JA. 2016. Oxygen consumption rate v. rate of energy utilization of fishes: a comparison and brief history of the two measurements. J Fish Biol 88:10–25. doi: 10.1111/jfb.12824

Peirson SN, Halford S, Foster RG. 2009. The evolution of irradiance detection: melanopsin and the non-visual opsins. Philos Trans R Soc Lond B Biol Sci 364:2849–2865. doi: 10.1098/rstb.2009.0050

Philipp DP, Cooke SJ, Claussen JE, Koppelman JB, Suski CD, Burkett DP. 2009. Selection for vulnerability to angling in largemouth bass. Trans Am Fish Soc 138:189–199. doi: 10.1577/t06-243.1

Polverino G, Santostefano F, Diaz-Gil C, Mehner T. 2018. Ecological conditions drive pace-of-life syndromes by shaping relationships between life history, physiology and behaviour in two populations of Eastern mosquitofish. Sci Rep 8. doi: 10.1038/s41598-018-33047-0

Quéméré E, Perrier C, Besnard AL, Evanno G, Baglinière JL, Guiguen Y, Launey S. 2014. An improved PCR-based method for faster sex determination in brown trout (*Salmo trutta)* and Atlantic salmon *(Salmo salar)*. Conserv Genet Resour 6:825–827. doi: 10.1007/s12686-014-0259-8

Réale D, Reader SM, Sol D, McDougall PT, Dingemanse NJ. 2007. Integrating animal temperament within ecology and evolution. Biol Rev 82: 291–318. doi:10.1111/j.1469-185X.2007.00010.x

Reddon AR. 2012. Parental effects on animal personality. Behav Ecol 23:242–245. doi: 10.1093/beheco/arr210

Redpath TD, Cooke SJ, Suski CD, Arlinghaus R, Couture P, Wahl DH, Philipp DP. 2010. The metabolic and biochemical basis of vulnerability to recreational angling after three generations of angling-induced selection in a teleost fish. Can J Fish Aquat Sci 67:1983–1992. doi: 10.1139/f10-120

Rosenfeld, J., Van Leeuwen, T., Richards, J. and Allen, D. 2015. Relationship between growth and standard metabolic rate: measurement artefacts and implications for habitat use and life-history adaptation in salmonids. J Anim Ecol, 84: 4–20. https://doi.org/10.1111/1365-2656.12260

Sbragaglia, V, Gliese, C, Bierbach, D, Honsey, AE, Uusi-Heikkilä, S, Arlinghaus, R. 2019. Size-selective harvesting fosters adaptations in mating behaviour and reproductive allocation, affecting sexual selection in fish. J Anim Ecol 88:1343–1354. https://doi.org/10.1111/1365-2656.13032

Schjolden J, Backström T, Pulman KGT, Pottinger TG, Winberg S. 2005. Divergence in behavioural responses to stress in two strains of rainbow trout *(Oncorhynchus mykiss)* with contrasting stress responsiveness. Horm Behav 48:537–544. doi: 10.1016/j.yhbeh.2005.04.008

Sharpe, D. M., Hendry, A. P. 2009. Life history change in commercially exploited fish stocks: an analysis of trends across studies. Evol appl 2:260–275. https://doi.org/10.1111/j.1752-4571.2009.00080.x

Sutter DAH, Suski CD, Philipp DP, Klefoth T, Wahl DH, Kersten P, Cooke SJ, Arlinghaus R. 2012. Recreational fishing selectively captures individuals with the highest fitness potential. Proc Natl Acad Sci USA 109:20960–20965. doi: 10.1073/pnas.1212536109

Svendsen MBS, Bushnell PG, Steffensen JF. 2016. Design and setup of intermittent-flow respirometry system for aquatic organisms. J Fish Biol 88:26–50. doi: 10.1111/jfb.12797

Therneau TM. 2018). coxme: Mixed Effects Cox Models. R package version 2.2-7. https://CRAN.R-project.org/package=coxme

Uusi-Heikkilä S, Whiteley AR, Kuparinen A, Matsumura S, Venturelli PA, Wolter C, Slate J, Primmer CR, Meinelt T, Killen SS, Bierbach D, Polverino G, Ludwig A, Arlinghaus R. 2015. The evolutionary legacy of size-selective harvesting extends from genes to populations. Evol Appl 8:597–620. doi: 10.1111/eva.12268

Uusi-Heikkilä S, Wolter C, Klefoth T, Arlinghaus R. 2008. A behavioral perspective on fishing-induced evolution. Trends Ecol Evol 23:419–421. doi: 10.1016/j.tree.2008.04.006

Väätäinen R, Huuskonen H, Hyvärinen P, Kekäläinen J, Kortet R, Arnedo MT, Vainikka A. 2018. Do metabolic traits, vulnerability to angling, or capture method explain boldness variation in Eurasian perch? Physiol Biochem Zool 91:1115–1128. doi: 10.1086/700434

Vainikka A, Tammela I, Hyvärinen P. 2016. Does boldness explain vulnerability to angling in Eurasian perch *Perca fluviatilis?* Curr Zool 62:109–115. doi: 10.1093/cz/zow003

Vilhunen, S., Hirvonen, H. 2003. Innate Antipredator Responses of Arctic Charr *(Salvelinus alpinus*) Depend on Predator Species and Their Diet. Behavioral Ecology and Sociobiology, 55, 1–10. www.jstor.org/stable/25063314

Vilhunen, S. 2006. Repeated antipredator conditioning: a pathway to habituation or to better avoidance? J Fish Biol, 68: 25–43. https://doi.org/10.1111/j.0022-1112.2006.00873.x

Vindas MA, Magnhagen C, Brännäs E, Øverli Ø, Winberg S, Nilsson J, Backström T. 2017. Brain cortisol receptor expression differs in Arctic charr displaying opposite coping styles. Physiol Behav 177:161–168. doi: 10.1016/j.physbeh.2017.04.024

Wickham H. 2009. ggplot2: Elegant Graphics for Data Analysis. Ggplot2: Elegant Graphics for Data Analysis:1–212. doi: 10.1007/978-0-387-98141-3

Wilson ADM, Brownscombe JW, Sullivan B, Jain-Schlaepfer S, Cooke SJ. 2015. Does Angling Technique Selectively Target Fishes Based on Their Behavioural Type? PLoS One 10. doi: 10.1371/journal.pone.0135848

Wong RY, Perrin F, Oxendine SE, Kezios ZD, Sawyer S, Zhou LR, Dereje S, Godwin J. 2012. Comparing behavioral responses across multiple assays of stress and anxiety in zebrafish (*Danio rerio*). Behaviour 149:1205–1240. doi: 10.1163/1568539x-00003018

Ylönen H, Kortet R, Myntti J, Vainikka A. 2007. Predator odor recognition and antipredatory response in fish: does the prey know the predator diel rhythm? Acta Oecol 31:1–7. doi: 10.1016/j.actao.2005.05.007

Ågren A, Vainikka A, Janhunen M, Hyvärinen P, Piironen J, Kortet R. 2019. Experimental crossbreeding reveals strain-specific variation in mortality, growth and personality in the brown trout *(Salmo trutta)*. Sci Rep 9. doi: 10.1038/s41598-018-35794-6

