## Appendix A for "Does parental angling selection affect the behavior or metabolism of brown trout parr?"

### Appendix A.
### Supplemental methods

#### Breeding and rearing of fish

Using the fully factorial breeding design, 27 families were produced in each of the four breeding groups. Two hundred eggs from each family were divided in two replicates, which were incubated in floating containers with nets as the bottom for water circulation (diameter ca. 10 cm). Containers were placed in 32 indoor fiber-glass tanks (0.4 $m^2$) and observed for egg mortality daily until hatching (no difference in mortality was observed between the groups). Soon after hatching, 25 alevins were haphazardly dip-netted from each family and pooled to two replicated matrix-specific groups of 225 individuals for each breeding group. When 25 fish were not available from a specific family, fish were taken equally from available families to sum 225 individuals in each tank. The two replicate groups were reared in dark green 0.4 $m^2$ -polyethylene hatchery tanks and fed with commercial fish food using automatic feeders, until the autumn of 2016.

#### Photoperiod manipulation

Fish from different breeding groups grown in the two replicate tanks were included nearly equally in each acclimation. The L:D group was divided in two groups, N = 30, in two tanks, and the 24h group was kept in a single tank, N = 40, due to lower number of individuals, until metabolic rate measurements started. Water temperature remained constant during the acclimation periods at mean 3.6 ± 0.1°C SD. The $O_2$ level remained constant at ~60% saturation (approx. 7.5 mg $O_2L^{-1}$) throughout the acclimation as the water originated from an ice-covered lake.

#### Sex determination using PCR

DNA samples were obtained for 82 out of 98 individuals. DNA was extracted from tail fin clip samples using a NucleoSpin DNA kit (Macherey-Nagel) following manufacturer's protocol, including an RNAse A treatment. For each individual, a positive control for successful DNA extraction was included in the PCR reaction (salmonid-specific 500 bp amplicon (Dash and Vasemagi 2014)). Duplex PCR reactions were prepared with Qiagen Multiplex Master mix, 0.2 µM primers and 2 µL template. Cycle conditions were: hot start 15 min at 95°C, then 36 cycles of denaturation at 94°C for 30 s, annealing at 56°C for 90 s, and elongation at 72°C for 90 s. The final extension was 72°C for 10 min. The PCR products were loaded on 2% agarose gel including SybrSafe, along with TrackIt 50 bp DNA ladder (ThermoFisher) and visualized using UV light.

#### Details of statistical analysis of behavior

The residuals of the initial model of *exploration intensity* were heteroscedastic, which was adjusted by using variance function $var(\epsilon_{ijk}) = \sigma^2\tilde{y}^{2\delta}$ to determine weights in function *lmer.* Parameter $\delta$ was first estimated using function *lme* from package *nlme* (Pinheiro et al. 2017) and used as fixed value $var(\epsilon_{ijk}) = 1.16^2\widehat{y_{ij}}^{2\times1.59}$ in the angling experiment, and $var(\epsilon_{ijk}) = 1.4^2\widehat{y_{ij}}^{2\times1.94}$ in burbot vs. control experiment. This approach satisfactorily modelled the heteroscedasticity.

The GLMM for *exploration tendency* was fitted with function *glmer* with the logit link function. The original model of angling selection experiment failed to converge with max|grad| = 0.0057, but it performed better than a converging model that did not include the random effect of arena using likelihood ratio test ($\chi^2$ = 13.523, df = 1, $P$ <0.001). No over-dispersion was found in the data.

For burbot vs control experiment, separate random intercepts were used for individuals in control and burbot groups. We additionally tested the homogeneity of variance in burbot and control conditions using Levene's test from (package *car*). For *exploration tendency*, a model with separate random intercepts for treatments was not significantly better than a model with a single intercept in a likelihood ratio test ($\chi^2$ =3.84, df = 2, $P$ = 0.15), and the former model also failed to converge. We therefore report the model with the single random intercept for exploration tendency.

#### Supplemental figures

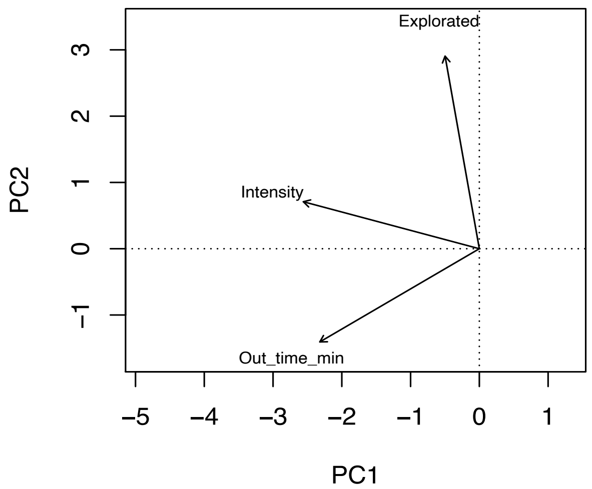


Fig. S1. Biplot showing the loading of different behavioral traits. Due to low eigenvalues and correlations among traits, each trait was analyzed in a separate univariate model.

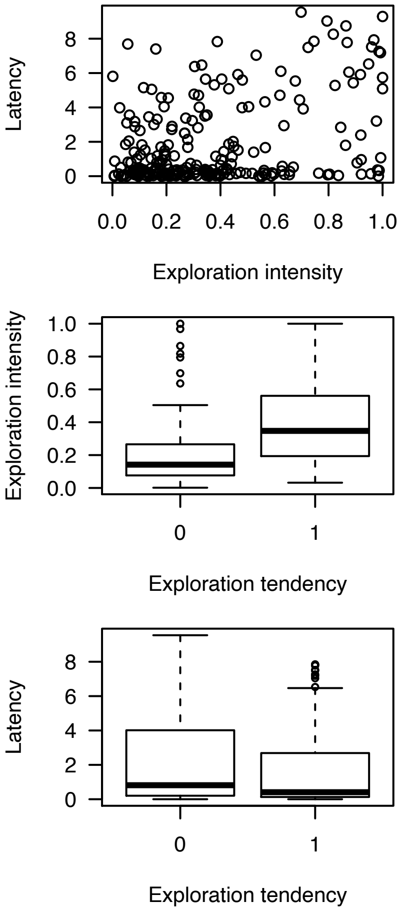


Fig. S2. Comparisons of behavioral variables in fish that emerged from start box during the trials.

**References for Supplemental Methods**

Dash M, Vasemagi A (2014) Proliferative kidney disease (PKD) agent Tetracapsuloides bryosalmonae in brown trout populations in Estonia. Diseases of Aquatic Organisms 109:139-148. doi: 10.3354/dao02731

Pinheiro J, Bates D, DebRoy S, Sarkar D and R Core Team (2017). nlme: Linear and Nonlinear Mixed Effects Models. R package version 3.1-131, https://CRAN.R-project.org/package=nlme
